## Supplementary material for "Spatial chromatin accessibility sequencing resolves high-order spatial interactions of epigenomic markers": QC_and_metrics

| sample | Base(Gb) | reads | Mean length | N50 | depth | Mapped reads | Order 1 reads num | Order 2 reads num | Order 3 reads num | Order 4-20 reads num | Order 21-50 reads num | Order >50 reads num | Total Fragments | Total contacts |
|---|---|---|---|---|---|---|---|---|---|---|---|---|---|---|
| nome7.5m_poreC_4T1 | 26.25 | 5,158,615 | 9,383 | 7,503 | 6.4 | 4,981,072 | 668,113 | 885,192 | 781,962 | 2,557,847 | 87,879 | 79 | 26,434,408 | 127,081,017 |
| nome15m_poreC_4T1 | 55.48 | 13,143,168 | 7,057 | 5,294 | 13.57 | 12,715,999 | 1,693,933 | 2,440,910 | 2,361,409 | 6,195,466 | 24,273 | 8 | 53,583,315 | 171,912,127 |
| nome3h_poreC_hek_wt (SCA_WT) | 129.94 | 44,985,375 | 2,888 | 4,446 | 36.92 | 43,188,859 | 9,224,815 | 9,575,918 | 6,955,451 | 17,299,368 | 133,093 | 214 | 169,906,655 | 492,502,643 |
| nome3h_poreC_hek_replicate_1 | 14.92 | 3,798,505 | 3,929 | 5,534 | 2.824 | 2,670,984 | 336,143 | 438,159 | 413,531 | 1,462,138 | 20,982 | 31 | 13,421,087 | 41,567,436 |
| nome3h_poreC_hek_replicate_2 | 20.32 | 5,179,923 | 3,923 | 5,563 | 3.826 | 3,639,666 | 461,053 | 581,799 | 559,610 | 2,004,178 | 32,974 | 52 | 18,549,210 | 56,692,566 |
| nome3h_poreC_hek_replicate_3 | 18.27 | 4,718,642 | 3,872 | 5,412 | 3.466 | 3,316,168 | 414,472 | 546,577 | 517,365 | 1,814,207 | 23,518 | 29 | 16,498,703 | 50,005,107 |
| poreC_hek_replicate_1 | 19.68 | 4,976,215 | 3,954 | 5,562 | 3.774 | 3,469,370 | 425,666 | 566,566 | 546,146 | 1,909,197 | 21,788 | 7 | 17,217,414 | 68,631,264 |
| poreC_hek_replicate_2 | 14.59 | 3,497,299 | 4,171 | 5,789 | 2.838 | 2,454,574 | 264,451 | 376,553 | 376,766 | 1,419,947 | 16,850 | 7 | 12,720,564 | 52,396,137 |
| poreC_hek_replicate_3 | 16.2 | 3,934,529 | 4,118 | 5,747 | 3.142 | 2,757,723 | 311,864 | 436,093 | 427,173 | 1,564,317 | 18,264 | 12 | 14,076,696 | 57,409,776 |
