## Supplementary material for "Spatial chromatin accessibility sequencing resolves high-order spatial interactions of epigenomic markers": data_table

| Sample | Assay | Target | Data accession id | Data link |
|---|---|---|---|---|
| hek293t_wild_type | Hi-C | 3D chromatin structure | GSE143465 | https://www.ncbi.nlm.nih.gov/geo/query/acc.cgi?acc=GSE143465 |
| hek293_wild_type | ATAC-seq | open chromatin | GSE108513 | https://www.ncbi.nlm.nih.gov/geo/query/acc.cgi?acc=GSE108513 |
| hek293t_wild_type | DNase-Seq | open chromatin | ENCSR000EJR | https://www.encodeproject.org/experiments/ENCSR000EJR/ |
| hek293t_wild_type | ChIP-Seq | CTCF | ENCSR135CRI | https://www.encodeproject.org/annotations/ENCSR135CRI/ |
| hek293t_wild_type | RNA-Seq | mRNA | GSE85161 | https://www.ncbi.nlm.nih.gov/geo/query/acc.cgi?acc=GSE85161 |
| hek293t_wild_type | pore-C | 3D chromatin structure | PRJNA917827 | https://www.ncbi.nlm.nih.gov/bioproject/?term=PRJNA917827 |
| hek293t_wild_type | SCA-Seq | 3D chromatin structure and accessibility | PRJNA917827 | https://www.ncbi.nlm.nih.gov/bioproject/?term=PRJNA917827 |
